# Age dependent changes in synaptic NMDA receptor composition in adult human cortical neurons

**DOI:** 10.1101/2020.01.21.913475

**Authors:** C.M. Pegasiou, A. Zolnourian, D. Gomez-Nicola, K. Deinhardt, J.A.R. Nicoll, A.I. Ahmed, G. Vajramani, P. Grundy, M.B. Verhoog, H.D. Mansvelder, V.H. Perry, D. Bulters, M. Vargas-Caballero

## Abstract

The molecular processes underlying the ageing-related decline in cognitive performance and memory observed in humans are poorly understood. Studies in rodents have shown a decrease in N-methyl-D-aspartate receptors (NMDARs) that contain the GluN2B subunit in ageing synapses, and this decrease is correlated with impaired memory functions. However, the age-dependent contribution of GluN2B containing receptors to synaptic transmission in human cortical synapses has not been previously studied. We investigated the synaptic contribution of GluN2A and GluN2B containing NMDARs in adult human neurons using fresh non-pathological temporal cortical tissue resected during neurosurgical procedures. The tissue we obtained fulfilled quality criteria by the absence of inflammation markers and proteomic degradation. We show an age-dependent decline in the NMDA/AMPA receptor ratio in adult human temporal cortical synapses. We demonstrate that GluN2B containing NMDA receptors contribute to synaptic responses in the adult human brain with a reduced contribution in older individuals. With previous evidence demonstrating the critical role of synaptic GluN2B in regulating synaptic strength and memory storage in mice, this progressive reduction of GluN2B in the human brain during ageing may underlie a molecular mechanism in the age-related decline in cognitive abilities and memory observed in humans.

## Introduction

Glutamate is the neurotransmitter of cortical and hippocampal pyramidal neurons and thus is a mediator cognitive functions. Glutamate-gated receptors of the NMDA subtype require intracellular depolarisation to relieve voltage-dependent extracellular Mg^2+^ block of the ion channel pore. Once opened, NMDAR channels can allow Ca^2+^ influx which triggers the synaptic plasticity processes thought to support many forms of learning and memory (Bliss and Collingridge, 1993; Takeuchi et al. 2013). NMDARs are tetrameric ion channels containing 2 GluN1 (obligatory) subunits and 2 GluN2 subunits, which in the adult forebrain most commonly comprise GluN2A and GluN2B. In addition to well characterised di-heteromers (GluN1/GluN2A or GluN1/GluN2B) (reviewed in Paoletti et al. 2013), tri-heteromeric forms (GluN1/GluN2A/GluN2B) have been selectively studied in expression systems (Hansen et al. 2014) and there is electrophysiological and biochemical evidence for NMDAR tri-heteromers and adult rodent brain (Al-Hallaq et al 2007; Rauner and Köhr, 2011; Stroebel et al. 2018). Furthermore, a short primate-specific GluN2A isoform (GluN2A-S) forms functional NMDAR together with GluN1 and accounts for one third of the total GluN2A protein in adult human cortex (Warming et al. 2019).

The subunit composition of NMDA receptors (NMDARs) determines their ion channel properties (Vicini et al. 1998), and protein-protein interactions with downstream molecular cascades that are linked to plasticity, survival and excitotoxicity (Hardingham and Bading, 2010; Lussier et al. 2015). Synaptic localisation of distinct NMDAR subunits is highly regulated by neurons and undergoes marked developmental changes in rodent neocortex with a switch from GluN2B to GluN2A rich synapses postnatally (Dumas, 2005; Mierau et al. 2004; Yashiro and Philpot, 2009). In adult synapses, a fraction of GluN2B-containing NMDARs is maintained in a highly regional-specific manner; sensory and association cortices maintain a lower proportion of GluN2B containing NMDARs than hippocampal and prefrontal cortex areas (Mierau et al. 2004; Wang et al. 2008; Kohl et al. 2011).

Previous research using human post-mortem tissue has shown that the ratio of *GRIN2B*/*GRIN2A* mRNA undergoes a reduction during early development similar to that found in many other species (Bar-Shira et al. 2015; Bagasrawala et al. 2016) and this is also evident at the protein level with an age-dependent reduction in GluN2B, and an increase in GluN2A, in the first years of life (Jantzie et al. 2013). Importantly, GluN2B has been found to co-assemble with PSD-95 in resected brain tissue from patients <17 years old (Ying et al. 2004), suggesting that GluN2B might contribute to synaptic transmission in young adult cortical synapses. However, it is unknown whether there is further reduction of GluN2B in adulthood or old age and whether GluN2B-containing NMDAR contribute to synaptic transmission in the adult brain.

Here, we tested directly whether GluN2B-containing NMDARs contribute to synaptic function in adult human cortical neurons. We used non-pathological tissue resected during neurosurgery from patients to test for NMDAR subunit composition in tissue homogenates and their association with the major synaptic scaffolding protein PSD-95. Furthermore we analysed synaptic transmission using patch clamp electrophysiology in brain slices of live temporal cortex. Our results show that NMDA/AMPA ratio decreases with age with a marked decline of GluN2B-NMDAR current in older synapses.

## Materials and Methods

### Tissue Collection

The use of human tissue complied with the Human Tissue Act (Southampton Research Biorepository study reference number: SRB002/14). Informed consent was obtained from all patients to use surgically-resected tissue not required for diagnostic purposes. Temporal cortex was chosen as its availability was sufficient to allow statistical analysis of data. Human brain slices were prepared as previously described by Verhoog et al. (2013). Briefly, resected tissue was obtained from temporal cortex of patients undergoing surgery for the removal of deeper structures. We obtained temporal cortical tissue from 17 individual neurosurgery cases (Table 1).

**Table 1.** Human tissue case data. F, female; M, male. Pathological tissue from patients 0011 and 0021 were used as positive control in Fig 1D. Cases that met exclusion criteria: cases 8 and 20 (Iba1 quantification), and case 11 (GluN2B cleavage). DNET, Dysembryoplastic neuroepithelial tumour.

| Case Number | Sex | Age | Diagnosis | Medication prescribed (time with medication) | Other medical history | Brain area |
| --- | --- | --- | --- | --- | --- | --- |
| 0001 | M | 49 | Hippocampal sclerosis | Carbamazepine (>1 year) | None known | Right temporal lobe |
| 0004 | M | 52 | Hippocampal sclerosis | Lamotrigine, Clobazam (>1 year) | None known | Right temporal lobe |
| 0005 | F | 60 | Hippocampal sclerosis | Levetiracetam, Phenytoin, Amitriptyline, Citalopram, Morphine, Omeprazole, Mebeverine (>1 year) | None known | Right anterior temporal lobe |
| 0007 | F | 21 | DNET | Lamotrigine, Eslicarbazepine (>1 year), Dexamethasone, Omeprazole | None known | Left anterior temporal lobe |
| 0008 | M | 71 | Glioblastoma | Phenytoin, Alfuzosin, Allopurinol, Dexamethasone, Paracetamol, Lansoprazole (<3 months) | None known | Right anterior temporal lobe |
| 0010 | M | 28 | Hippocampal sclerosis | Lamotrigine, Clobazam, Carbamazepine (>1 year) | None known | Right posterior lateral temporal lobe (inferior temporal gyrus) |
| 0011 | F | 42 | Glioma | N/A | None known | Left anterior temporal cortex |
| 0014 | M | 32 | Hippocampal sclerosis | Lamotrigine, Levetiracetam, Pregabalin, Citalopram, Paracetamol, Salbutamol (>1 year) | Asthma, Depression | Right anterior temporal lobe |
| 0016 | F | 36 | Hippocampal sclerosis | Gabapentin, Lamotrigine, Citalopram, Senna, Omeprazole, Paracetamol (>1 year) | None known | Left anterior temporal lobe |
| 0017 | F | 62 | Hippocampal sclerosis | Lamotrigine, Carbamazepine, , Diazepam, Pregabalin, Citalopram Prochlorperazine (>1 year) | Depression, Hypertension, Gastroreflux disease | Right anterior temporal lobe |
| 0018 | M | 30 | Cavernous malformation | Lamotrigine, Levetiracetam (>1 year) | None known | Right anterior temporal lobe |
| 0020 | F | 70 | Arteriovenous malformation | Fentanyl Patch, Paracetamol, Amlodipine, Omeprazole, Estradiol, Avastatin, Qvar 100, Colecalciferol, Prednisolone, Nitrofurantoin (<1 month) | Hysterectomy, Urostomy, Acute Myocardial Infarction, Hypertension, Polymyalgia Rheumatica, Hypercholesterolaemia, Recurrent Urinary Tract Infections | Right lateral temporal lobe (middle temporal gyrus) |
| 0021 | M | 49 | Hippocampal sclerosis | Carbamazepine, Atenolol (>1 year) | Glandular Fever | Left posterior temporal lobe |
| 0022 | F | 58 | Intracerebral and subarachnoid haemorrhage | Dihydrocodeine, Nimodipine, Paracetamol (N/A) | Multiple Aneurysms | Right anterior temporal lobe |
| 0024 | F | 50 | Cavernous malformation | Lamotrigine, Ibuprofen, Paracetamol (>1 year) | Gall stones, Hysterectomy | Right lateral temporal lobe (superior temporal gyrus) |
| 0026 | M | 27 | Mesial temporal DNET with signal changes in the hippocampus | Carbamazepine, Clobazam, Levetiracetam, Zonisamide, Morphine, Dihydrocodeine, Dexamethasone, Omeprazole, Paracetamol, Colecalciferol (>1 year) | None known | Right anterior temporal lobe |
| 0028 | F | 38 | Epilepsy | Zopiclone, Amitriptyline, Zonisamide, Citalopram, Zafirlukast, Cinnarizine, Loestrin (>1 year), Senna | Asthma | Right anterior temporal lobe |

In all cases, resected neocortical tissue was located well outside the epileptic focus or tumour and displayed no structural abnormalities in preoperative magnetic resonance imaging investigations (Fig 1A). The tissue collected was identified as macroscopically normal at the time of collection by the operating team. The tissue was immediately placed in ice-cold artificial cerebrospinal fluid (ACSF) solution while still in theatre. ACSF contained the following (in mM): 110 choline chloride; 26 NaHCO3; 10 D-glucose; 11.6 sodium ascorbate; 7 MgCl2; 3.1 Sodium pyruvate; 2.5 KCl; 1.25 NaH2PO4.2H2O; 0.5 CaCl2. Tissue was then taken to the laboratory and fresh frozen for biochemistry, fixed for immunohistochemistry or processed for slice electrophysiology. Transfer time between operating theatre and laboratory was approximately of 10-15 minutes.

**Figure 1.**
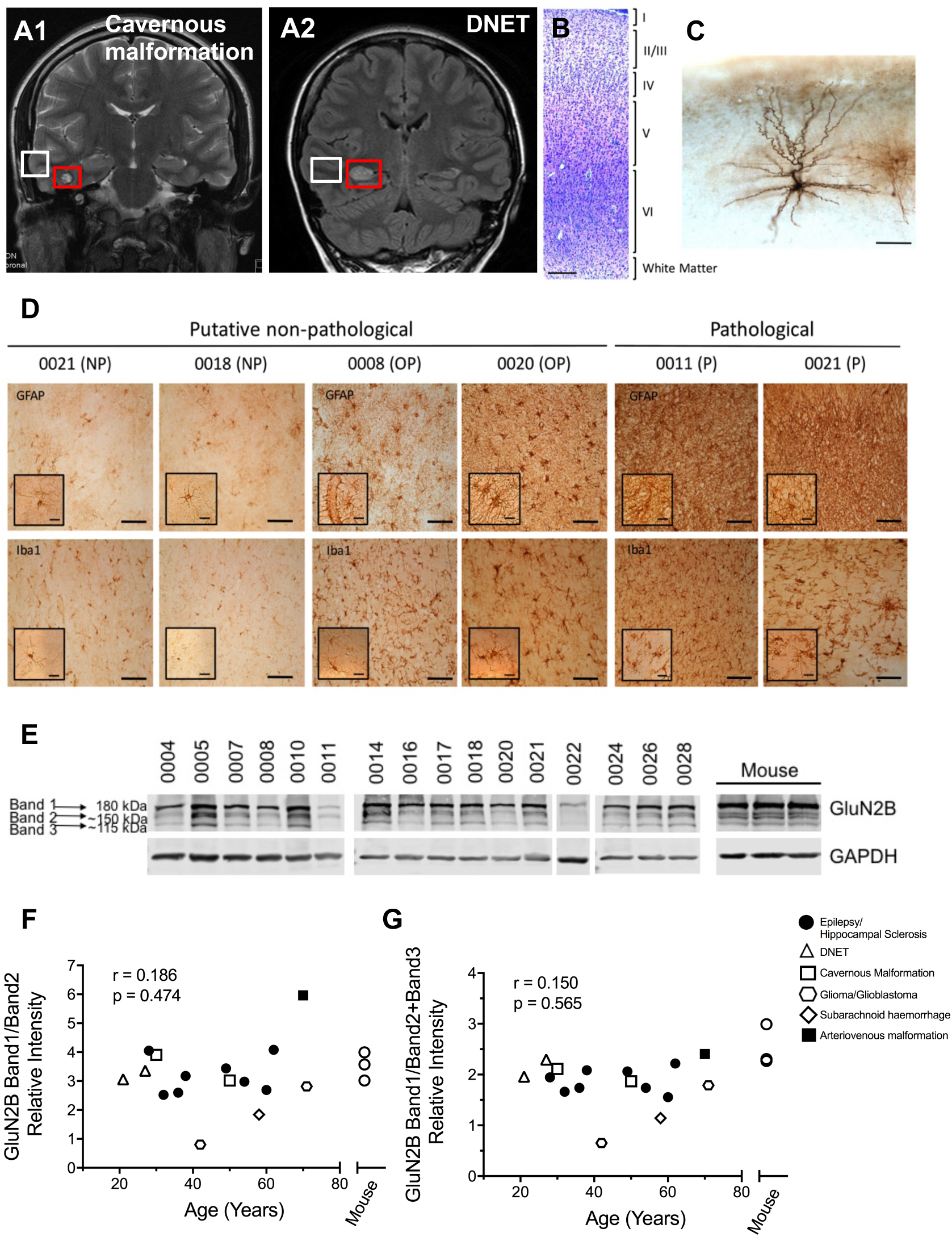
Characterisation of human brain tissue resected from neurosurgical cases. (A) MRI images from two patients. Coronal T2 image of a cavernous malformation (A1) and coronal FLAIR image of a DNET (A2) taken prior to surgery. Red box: area of pathology necessitating neurosurgical removal, white box: resected tissue used for analysis. (B) Nissl stain in fixed tissue showing the preservation of cortical layer architecture. (C) Visualisation of a LII-III pyramidal neuron that was recorded in patch clamp mode and dialyzed with biocytin, fixed, and DAB stained. (D) Immunohistochemical analysis of putative non-pathological resected tissue compared with pathological tissue resected from the region of underlying pathology. Representative images of immunostaining for inflammatory markers, GFAP and Iba1, of putative non-pathological (PNP) resected temporal cortex showing no pathology (NP), of putative non-pathological resected temporal cortex where we observed pathology (OP), and of known pathological tissue, resected from the regions of underlying pathology (P). (E) GluN2B immunoblot using resected human samples; lysates made from fresh-frozen human temporal cortices tissue and from fresh-frozen mouse temporal cortices (n=3). (F) Band1/Band2 and (G) Band1/(Band2+Band3) ratio plotted against age. No correlation observed between the ratios and subject age (Spearman’s coefficient of correlation (r) and p-value (p) are indicated). Scale bars in (**B**) = 200 μm, in (**C**) = 100 μm, in (**D**) = 100 μm (Inset = 20 μm). Symbols in F and G correspond to the neurological condition necessitating neurosurgery.

Animal care and experimental procedures were conducted in accordance with UK Home Office regulations under the Animals (Scientific Procedures) Act of 1986. Mice were decapitated following isoflurane anaesthesia. Brains were extracted in ice-cold ACSF and sliced or snap-frozen. All brain tissue samples were stored in the -80°C freezer until lysed.

### Acute Brain Slice Preparation

Brain slices (350 μm thick) were prepared in ice-cold choline ACSF solution (Verhoog et al. 2013) using a Campden Instruments vibrating microtome and kept for 30 min at 35-37°C, then at room temperature in recording ACSF containing the following (in mM): 126 NaCl; 2 CaCl2; 10 glucose; 2 MgSO4.7H2O; 3 KCl; 1.25 NaH2PO4.2H2O; 26.4 NaHCO3; pH 7.2-7.4. The ACSF was bubbled with carbogen gas (95% O2/5% CO2) and had an osmolarity of ∼300 mOsm.

### Electrophysiological Recordings

Following previously published methods (Verhoog et al. 2013), slices were transferred to a submerged-style recording chamber and superfused with recording ACSF at a rate of ∼1.5ml/min. Whole-cell voltage clamp recordings were performed using glass pipettes (4-6 MΩ) pulled from borosilicate glass, yielding a series resistance of 10-20 MΩ. Recordings were made at room temperature (21-25°C) using Cs-gluconate-based intracellular solution, containing (in mM): 70 Gluconic acid; 10 CsCl; 5 NaCl; 10 BAPTA free acid; 10 Hepes; 10 QX-314; 0.3 GTP; 4 Mg-ATP; pH was titrated to 7.25 with CsOH. The estimated final Cs concentration for the intracellular solution was 120 mM. The final osmolarity was 280 ± 5 mOsmol-1. All voltage values were corrected for the liquid junction potential of -15 mV which was measured directly. Biocytin (2mg/ml) was added for labelling (Fig 1C) and following recordings, the slice was fixed in 4% paraformaldehyde for 24 hours and stored in PB. Slices were washed in 0.1M PB, dehydrated using increasing ethanol percentages (70%, 80%, 95% and 2×100%, for 5 minutes each) and then processed using an avidin–biotin–peroxidase kit (Vector laboratories).

Excitatory postsynaptic currents (EPSCs) were evoked using a stimulus isolator unit (ISO-Flex, A.M.P.I.) connected to a monopolar extracellular stainless-steel electrode, which was placed near an apical dendrite and within 100-150 µm from a LII-III pyramidal neuron soma. Stimulation strength was adjusted to yield a ∼200 pA EPSC amplitude at a holding potential of −70 mV to minimise space-clamp error. Synaptic stimulation was evoked at 0.07 Hz with 100 µs stimulus length. Series resistance was not compensated during recordings but was monitored before each stimulation with a 5 mV 50 ms step pulse. Recordings were terminated if series resistance changed by more than 20%. Data were low-pass filtered at 2 kHz and acquired at 20 kHz with an Axon Multiclamp 700B amplifier (Molecular Devices, Sunnyvale, U.S.A.) using MATLAB (Mathworks, Natick, U.S.A.) and custom software (MatDAQ, Hugh P.C. Robinson, University of Cambridge 1995-2013). EPSCs were recorded 10 minutes after initiating a whole-cell patch to allow the dialysis of the Cs-based internal solution. EPSCs were recorded in the presence of gabazine (also known as SR-95531, Tocris Bioscience) at a concentration of 3 μM to measure NMDAR currents in the absence of GABA_A_-mediated currents (Vargas-Caballero et al. 2011). To measure the contribution of AMPAR and NMDAR-mediated currents to individual EPSCs, the holding membrane potential was clamped at −70mV and +40mV, respectively, in alternating sweeps. Cells were maintained at −70 mV in between stimuli and the membrane potential was changed to holding potential 2 sec prior to synaptic stimulation. Evoked EPSCs were analysed using custom-written MATLAB scripts. A minimum of 5 continuously recorded sweeps for each AMPAR and NMDAR were measured. AMPAR current was measured at its peak and NMDAR current was measured as the average current at 55-57ms following stimulation, at which time the AMPAR showed decay to ∼5% of its peak amplitude. The NMDAR/AMPAR ratio was calculated by dividing the amplitude of NMDAR current measured by the peak AMPAR current. Successful patch clamp recordings in whole-cell configuration could be made up until 12 hours after resection using this tissue.

Averaged evoked NMDAR currents (from peak back to baseline) were fitted with least squares (Matlab) using a double exponential equation as follows:

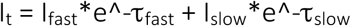

where I_fast_ and I_slow_ were the amplitudes of the fast and slow decay components, and τ_fast_ and τ_slow_ are the decay time constants. Following Stocca and Vicini (1998) we used a weighted mean decay time constant to compare a single experimental value across conditions:

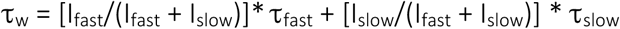

A single τ_w_ value was obtained per patient by averaging τ_w_ from different cells.

### Co-immunoprecipitation

Fresh frozen human tissue blocks immediately adjacent to sliced/fixed tissue, were stored at - 80°C after resection was homogenized on ice using a Pellet Pestle in SDS-free RIPA buffer (2x Phosphate Buffered saline (PBS); 1% sodium deoxycholate; 1% NP40, 5mM EDTA; Halt protease and phosphatase inhibitor cocktail (ThermoFisher Scientific). Homogenates were centrifuged at 14,000 rpm for 10 min 4°C and supernatants were collected for BCA assaying (ThermoFisher Scientific) to determine protein concentrations. One lysate sample for each human case was set aside for Western blotting as input lane. For immunoprecipitation, a total of 1mg of protein was diluted in 1ml of RIPA buffer. A pre-clear step was performed by adding protein A-agarose beads (Merck Millipore) to the lysates, while rotating for 30min at 4°C. Following centrifugation at 10,000 rpm for 2 min, the supernatants were collected and samples were incubated overnight at 4°C with PSD-95 antibody raised in rabbit (CST #2707; 1:50 dilution), or rabbit IgG as control. Samples were precipitated by incubating with protein A-agarose beads for 1 hr at 4°C. Precipitated proteins were released from the beads by heating at 95°C for 5 min in 4X loading sample buffer prior to SDS-PAGE along with 18.75 μg of input.

### SDS-PAGE and Western Blotting

Equal amounts of protein (28μg) were separated in 7.5% acrylamide gels by SDS-PAGE and transferred onto nitrocellulose membranes. Membranes were blocked in 5% (w/v) non-fat milk for 1hr at room temperature and incubated overnight at 4°C in 5% (w/v) bovine serum albumin (BSA) containing 0.1% (v/v) Tween-20 and one of the following primary antibodies: anti-NMDAR2A (ab133265; 1:1000; Abcam); anti-NMDAR2B (610416; 1:1000; BD Neurosciences), anti-PSD-95 (D27E11; 1:1000; CST), and GAPDH (D16H11; 1:1000; CST). Membranes were washed 3 times with Tris-buffered saline (TBS) containing 0.1% Tween-20 (TBS-T) and probed with fluorophore-conjugated goat anti-mouse/-rabbit secondary antibody (1:10000; LI-COR). Proteins were visualised using the Odyssey infrared scanner (LI-COR) and bands were quantified as a proportion of housekeeping protein GAPDH using Image Studio Light Software.

### Immunohistochemistry

Tissue was immediately fixed in 4% paraformaldehyde for 24 hrs and cut into 35 µm free-floating sections for immunohistochemistry. All washes were performed with PBS containing 0.1% Tween-20 (PBS-T 0.1%), unless stated otherwise. Sections were incubated in 0.3% H2O2, 10% methanol in PBS-T 0.1% for 30mins and then in blocking solution containing 5% goat serum (Sigma Aldrich), 5% BSA (Fisher Scientific, BP1600) in PBS-T 0.2% for 1hr. Sections were then incubated overnight at 4°C in blocking solution containing one of the following primary antibodies: anti-GFAP (1:2000; Merck Millipore) and anti-Iba1 (1:500; Wako). Sections were then washed (3 x 5min) and incubated with the appropriate biotinylated secondary antibody (1:200, Biotinylated Goat Anti-Mouse Antibody (BA-9200), 1:200, Biotinylated Goat Anti-Rabbit Antibody (BA-1000), Vector Labs). Sections were then incubated in the avidin-biotinylated horseradish peroxidase complex (ABC) for 30 minutes. Following the ABC incubation sections were mounted onto gelatinised slides and were developed using 3,3’-diaminobenzidine (DAB) precipitation. Sections were washed (3 x 5min) in 0.1M PB, dehydrated using increasing ethanol percentages (70%, 80%, 95% and 2×100%, for 5 minutes each). Sections were incubated in xylene (2×10minutes each) and mounted using Entellan^®^ New, mounting medium. Images from stained sections were taken using a Leica Microscope (Leica DM5000B) with Q software for image capture.

Sections were visualised using 3,3’-diaminobenzidine (DAB) precipitation under a light microscope (Leica DM5000B) with Q software for image capture. GFAP- and Iba1-positive cells were counted in 12 layer II-III sampling fields per case (chosen at random from 2-3 different slices) of temporal cortical tissue from putative non-pathological and pathological (sclerotic hippocampus or glioma) samples with available tissue using ImageJ (NIH) software.

## Results

We used non-pathological cortical tissue samples resected during neurosurgical procedures to obtain access to deep brain lesions (Fig. 1A, Table 1). The tissue was collected in theatre and rapidly processed using the brain slice technique (Verhoog et al. 2013). The well-preserved cortical architecture of these slices (Fig. 1B) allowed for electrophysiological recordings in putative LII-III pyramidal neurons that were identified by their location within cortical layers and their morphology (Fig. 1C). First, we characterised the quality and health of collected tissues using fixed or snap-frozen samples of brain tissue that were immediately adjacent to that used for electrophysiology. Reactive astrocytes and activated microglia are reliable inflammatory indicators of disease in mammalian brain tissue (Perry et al. 2010) (Figure 1D). As positive controls for inflammatory pathology, we stained pathological sections (epileptic foci N = 2 patients, tumour tissue N = 1 or cavernous malformation tissue N = 1) with GFAP to label astrocytes and Iba1 to label microglia. We compared putative non-pathological tissue (Supplementary Fig. 1A-E) against pathological tissue (cases 11 and 21). Qualitatively, the putative non-pathological tissue showed normal GFAP-positive cell morphology in contrast with highly ramified and fibrotic astrocytes observed in pathological tissue. Furthermore, the GFAP density was significantly lower in putative non-pathological tissue than in pathological tissue. In our Iba1 analyses, we found two cases with activated microglia density similar to that of positive controls (Fig. 1D and Suppl. Fig. 1C). We therefore labelled these two cases as observed pathology (OP) and excluded them from synaptic analyses. The density of microglia in the human brain does not change with age, as recently reported by us (Askew et al. 2017). Here, we observed densities in accordance with previously published data, and similarly did not observe significant age-dependent changes in astrocyte density (r = -0.03 p = 0.92) or microglia density (r = -0.54 p = 0.09).

Next, we sought to further control for resected tissue quality by measuring the ratio of full length to cleaved GluN2B protein, which has previously been demonstrated as a robust control for synaptic proteome integrity in post-mortem human brain tissue (Bayés et al. 2014). Using Western blot analysis of GluN2B in fresh frozen samples with an antibody that recognises full-length protein (Band 1, 180kDa) and cleaved products (Bands 2 and 3, 150 and 115kDa), we measured the ratio of full-length to cleaved GluN2B (Figure 1E-G). All except one of our samples met the criteria of Band 1/Band 2 ratio >1 (Bayés et al. 2014) demonstrating lack of proteome degradation. Given the observed degradation of full-length GluN2B, the sample showing a ratio of <1 (case 0011), was excluded from synaptic analyses. All fresh-frozen tissue samples showed the presence of GluN2B cleavage products, likely indicating functional and turnover-related regulation of the channel by proteases such as calpain (Wu et al. 2005). The full-length-to-cleaved band ratio was not altered with age (Figure 1F-G). We observed a similar GluN2B protein profile in rapidly processed adult mouse brain tissue (Fig 1E-G), which confirmed that basal cleavage of GluN2B in fresh frozen tissue was not an artefact of tissue collection. This shows that high quality tissue can be obtained from human patients irrespective of age.

To test whether total protein levels decrease with age, we performed further Western blot quantifications of GluN2A and GluN2B subunits. We found that both GluN2A and GluN2B protein levels decreased with age, and a trend (p = 0.08) was observed for GluN2A-S (Fig 2A-D). Using PSD-95 pull-down from brain homogenates we were able to observe both GluN2A and GluN2B in samples from 27, 49 and 50 year old patients, however in a sample from a 58 year old we were not able to see GluN2B co-immunoprecipitated with PSD-95 (Fig 2E). Not all samples yielded successful precipitated PSD-95 lanes and thus we did not quantify co-precipitation. To analyse whether the observed decrease in GluN2A and GluN2B NMDAR subunits was associated with overall reduction in synaptic proteins we also tested available samples for synapsin (pre-synaptic), PSD-95 (post-synaptic) and the AMPAR subunit GluA1 (Fig 3A). We observed that there was no significant correlation with age in the amount of these proteins in cortical homogenate normalised against tubulin (Fig 3B-G) or against total protein load (Ponceau stain of Western blot membrane, data not shown). However, a significant negative correlation of GluN2B protein levels with age was consistently observed when normalising against tubulin (Fig 3B), Ponceu stain (not shown) or against PSD-95 (Fig 3F) as a postsynaptic protein control.

**Figure 2.**
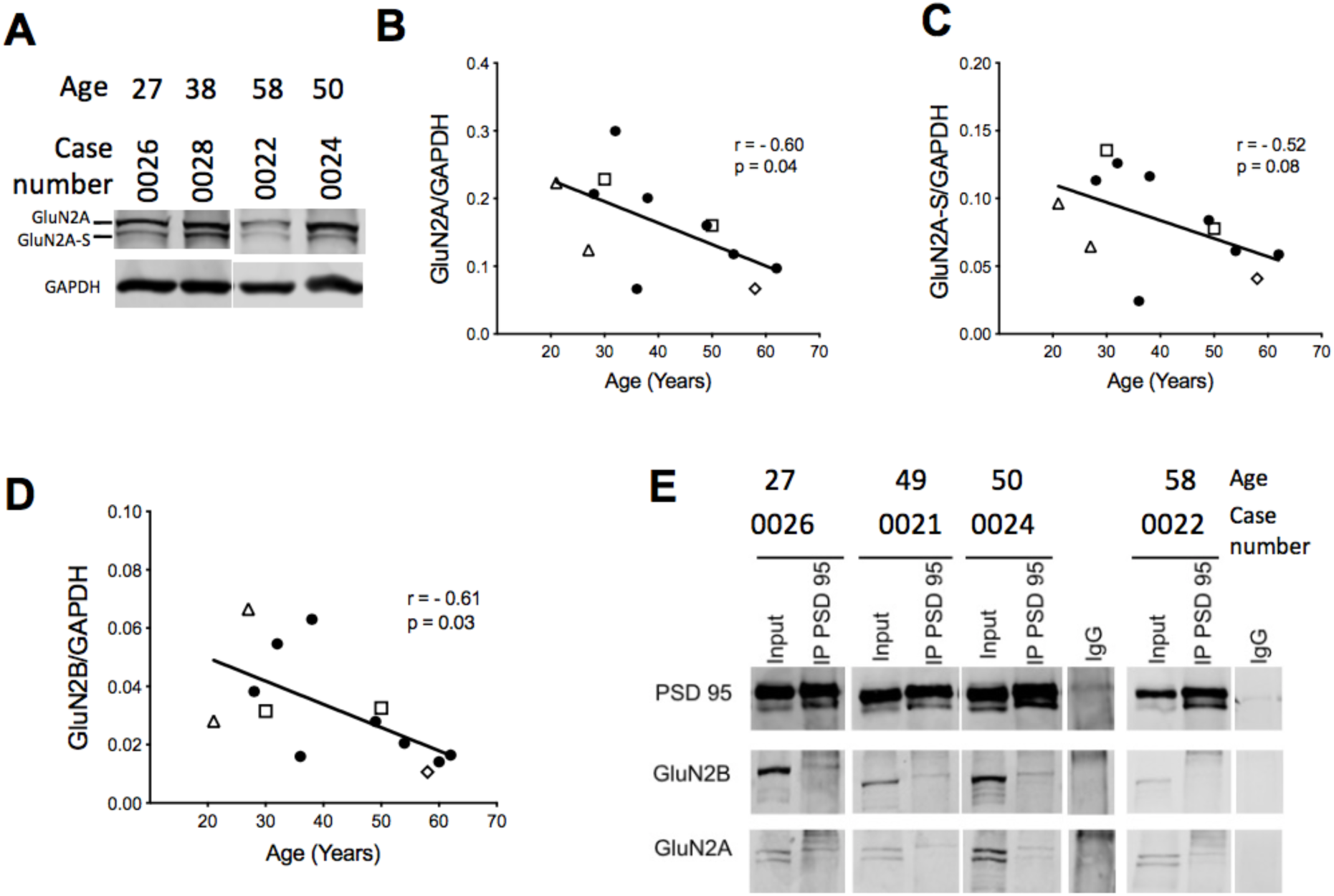
GluN2A and GluN2B protein levels in adult human temporal cortical tissue homogenate show an age-dependent decline. **(A) Representative examples of GluN2A blot showing GluN2A and GluN2A-S bands.** Quantification for GluN2A **(B)** and GluN2A-S (C), and GluN2B (B, from blots shown in Fig. 1E) relative to the constitutive protein GAPDH. Spearman’s coefficient of correlation (r) and p-value (p) are indicated for each measure. (E) Representative blots of GluN2A and GluN2B subunits co-immunoprecipitated with PSD-95 using human temporal cortical tissue lysates. Symbols in B, C and D correspond to the neurological condition necessitating neurosurgery as depicted in Figure 1.

**Figure 3.**
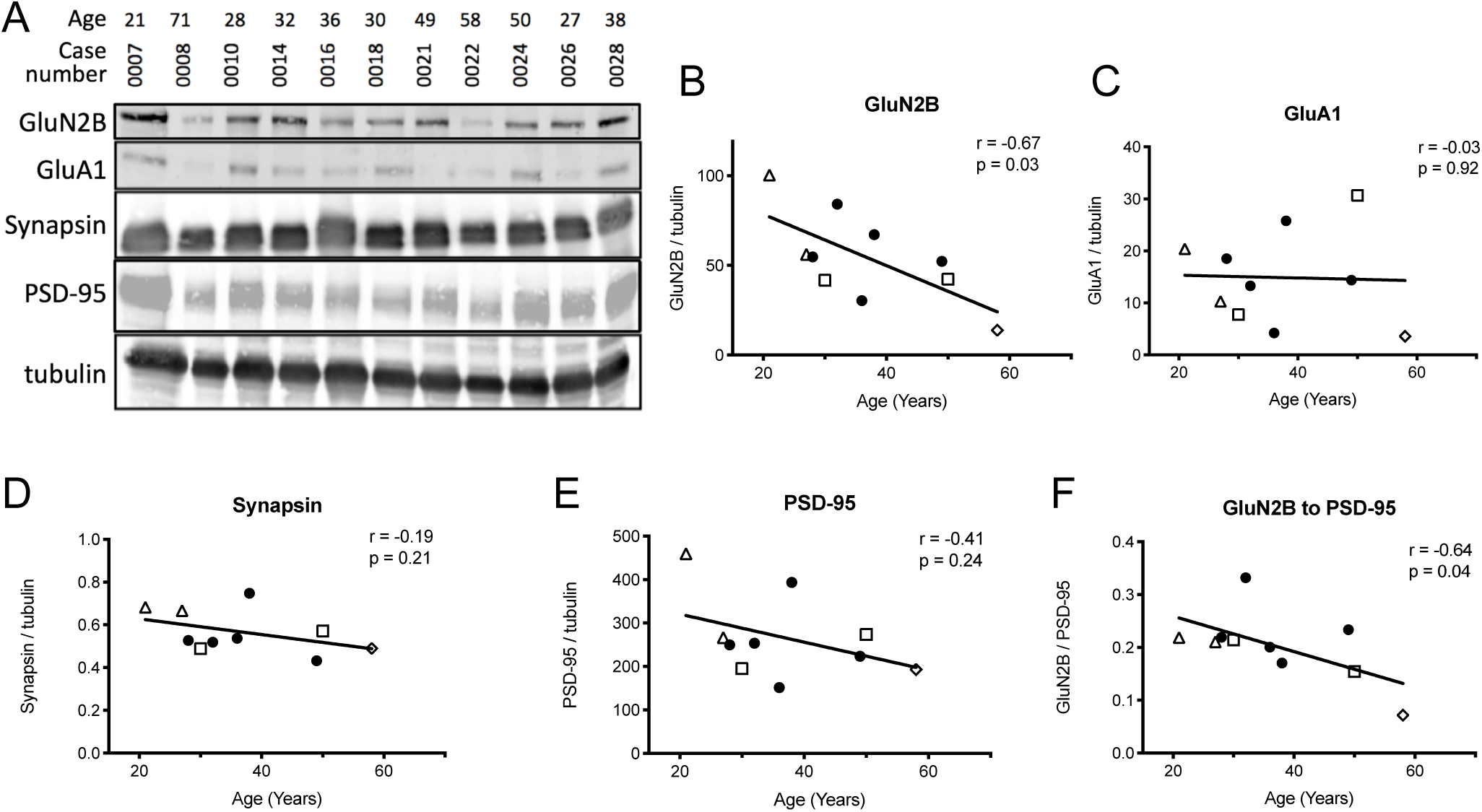
Age dependence of GluN2B and other synaptic protein expression in human temporal tissue homogenate. (A) Western blot for GluN2B and other synaptic proteins with tubulin as loading control. Quantification of GluN2B (B), GluA1 (C), synapsin (D) and PSD-95 (E) normalised to tubulin and plotted against age. (F) Quantification of GluN2B normalised to PSD-95 (G) and plotted against age. Spearman’s coefficient of correlation (r) and p-value (p) are indicated for each measure. Symbols in C-G correspond to the neurological condition necessitating neurosurgery as depicted in Figure 1. Case 0008 in (A) was pathological and was not quantified.

We next analysed patch clamp recordings to determine whether these receptors participate in basal synaptic transmission. NMDA and AMPA currents exhibited current voltage relationships similar to those observed in rodents (Figure 4A-B). We analysed the age-dependence of NMDA/AMPA receptor ratio by measuring AMPAR component at -70 mV and NMDAR component at sustained +50 mV depolarisation to relieve the Mg^2+^ block (Mierau et al. 2004; Vargas-Caballero and Robinson, 2004), in subsets of cells with and without pre-incubation with the selective GluN2B inhibitor, Ro 25-6981 (500 nM), which shows similar pharmacological properties in both rat and human recombinant receptors (Hedegaard et al. 2013). The NMDA/AMPA ratio showed a significant decrease with age (Figure 4C), suggesting an age-dependent reduced contribution of NMDARs to synaptic transmission. In contrast, the NMDA/AMPA ratio remained constant across ages in recordings from slices pre-incubated with Ro 25-6981 (Figure 4D), which at 500 nM inhibits recombinant NMDAR di-heteromers (GluN1/GluN2B). This suggests that GluN2B loss could be a key determinant of reduced synaptic NMDAR input to ageing temporal cortical synapses.

**Figure 4.**
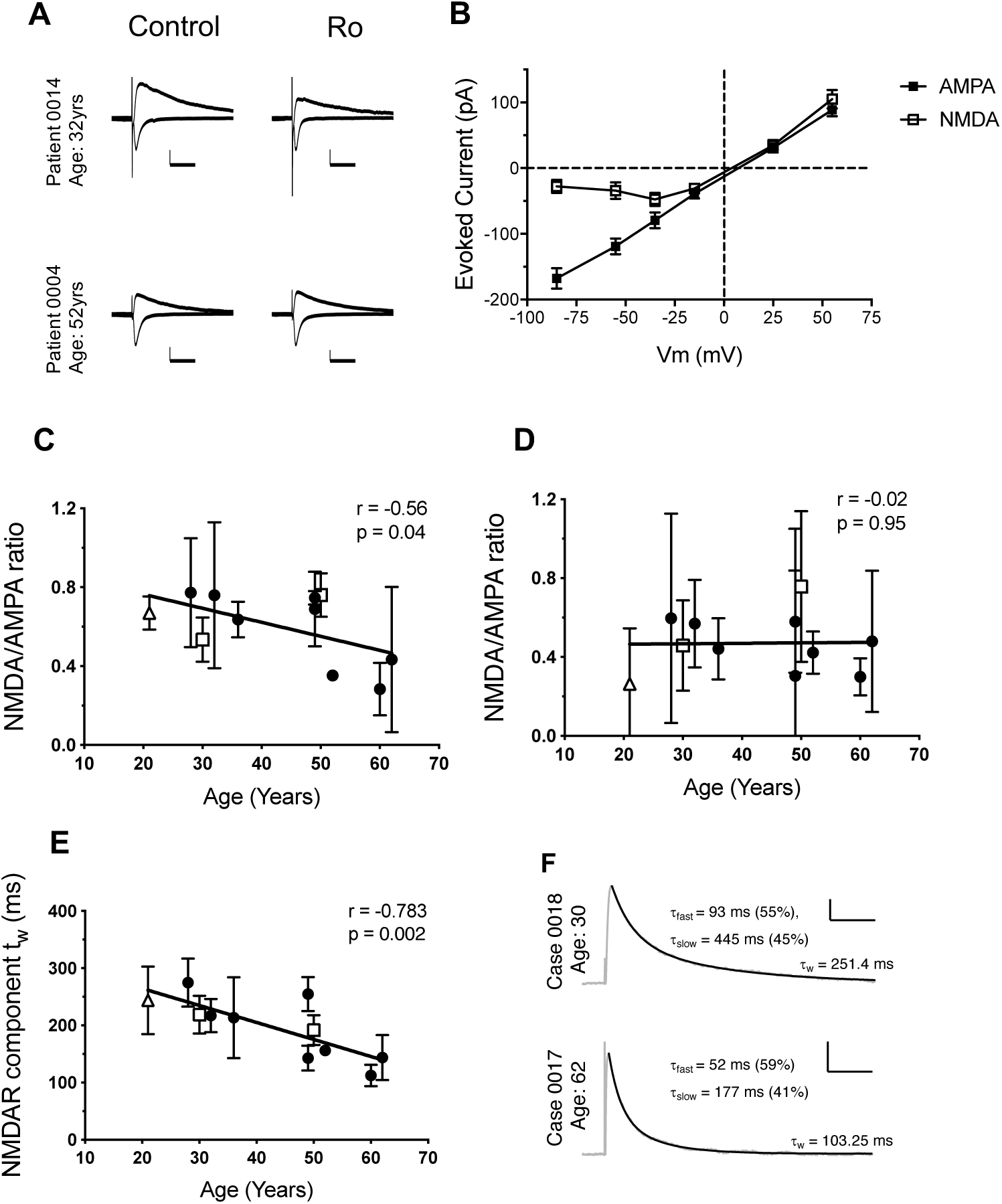
NMDA/AMPA responses from human cortical synapses with or without specific GluN2B inhibitor and their age dependence. (A) Representative NMDA- and AMPA-mediated currents from adult human brain slices in LII-III temporal cortex voltage clamped at -70 and +50mV with and without 500 nM Ro 25-6981 treatment. NMDAR-mediated currents from a 32-year-old are slow-decaying and with Ro treatment they producing a smaller and faster decaying outward current. NMDAR-mediated currents from a 52-year-old show no sensitivity to Ro 25-6981 treatment. (B) Average I/V relations for NMDA and AMPA responses (n=7 patients). I/V relations for NMDA-mediated responses show a voltage dependence (characteristic ‘J-shaped’ curve), showing the block of the receptor at hyperpolarized membrane potentials. I/V relations for AMPA-mediated responses are linear. (C) NMDA/AMPA ratios are negatively correlated with age under control conditions. (D) No correlation was observed between NMDA/AMPA ratios and age for recordings carried in 500 nM Ro 25-6981. (E) The weighted NMDAR time constant τ_w_ shows an age dependent decline, suggesting a reduced contribution of the slow decaying GluN1/GluN2B containing NMDARs. Each data point represents the average measure ± SEM per patient. (F) Sample lines of best fit. Scale bars in (A) = 50 ms, 50 pA and (F) = 200 ms, 50 pA. Scatter dot plots with error bars in graphs represent mean ± SEM per patient. Spearman’s coefficient of correlation (r) and p-value (p) are indicated for each measure. Symbols in C, D and E correspond to the neurological condition necessitating neurosurgery as depicted in Figure 1.

In a subset of experiments, NMDAR currents were measured before and after application of Ro 25-6981 in the same cell. These experiments required holding neurons in whole-cell configuration for >30 minutes to allow time for initial dialysis of intracellular voltage-clamp solution, acquisition of control data, wash in of bath-applied Ro and, importantly, to allow time for full NMDAR current inhibition by the use-dependent antagonist Ro 25-6981 (Fischer et al. 1997). We observed that NMDA/AMPA ratios from subjects younger than 45 showed sensitivity to Ro treatment while those for older than 45 subjects did not (<45 years old NMDA/AMPA ratio before = 0.88 ± 0.20, after Ro 25-6981 treatment = 0.46 ± 0.05 N = 3, and >45 years old NMDA/AMPA ratio before = 0.53 ± 0.15 after Ro 25-6981 treatment 0.474 ± 0.08, p < 0.05, N = 5). These data also show that the fraction of NMDAR current inhibited by Ro was larger in synapses from younger individuals (42% ± 0.15 versus 1% ± 0.16 in older individuals, p < 0.05). These findings are consistent with our results above using Ro preincubation.

Given that recombinant NMDARs containing either GluN1/GluN2A and GluN1/GluN2B have distinct postsynaptic current decay kinetics (Stocca and Vicini, 1998), we fitted a double exponential decay function to all recorded currents and obtained a weighted time constant value (τ_w_, see methods) for each case. We found that τ_w_ decreased significantly with age, this is consistent with synapses from younger individuals having a higher proportion of synaptic GluN2B containing receptors that is progressively reduced with age (Fig 4E). Interestingly, we found that in the presence of Ro 25-6981 500 nM there was still significant decrease of τ_w_ with age albeit at lower significance value (p< 0.05) and with a smaller correlation coefficient than in the absence of Ro 25-6981 (r = -0.57). It is possible that tri-heteromers, that have τ_w_ larger than GluN1/GluN2A but lower than GluN1/GluN2B di-heteromers are the cause of this enhanced τ_w_ in younger individuals. In this instance, Ro 25-6981 (an ifenprodil derivative) would only achieve partial inhibition of tri-heteromers even at saturating antagonist concentrations (Stroebel et al. 2018). However, we cannot rule out whether other slower-decaying NMDAR variants (e.g. GluN2D or those containing GluN1-1a) are present at a younger age in human cortical synapses.

## Discussion

A sharp developmental increase in GluN2A and a decrease in GluN2B-containing NMDARs has been observed in many model systems including rodents (Dumas, 2005; Yashiro and Philpot, 2009 and reviewed in Paoletti et al. 2013). In humans, an 8% reduction in *GRIN2B* mRNA expression during mid-gestation was accompanied by a 21% increase in *GRIN2A* mRNA expression (Bagasrawala et al. 2016), these observations are consistent with larger scale mRNA analyses across the human lifespan showing that, while minor changes occur with GRIN2B during embryonic development and adult life, a major increase in GRIN2A expression is observed during gestation (Bar-Shira et al. 2015).

Although GluN2A containing channels are broadly considered the major carriers of NMDAR-mediated synaptic current in the adult forebrain (Hildebrand et al. 2014), adult GluN2B-containing NMDARs are required in adult brain circuits to regulate synaptic strength and memory in mice. In hippocampal CA3-CA1 synapses the levels of GluN2B expression in CA3-CA1 synapses are correlated with the ability of synapses to undergo long term potentiation (Kohl et al. 2011). The high affinity association between GluN2B subunits with CaMKIIα (Barria and Malinow 2005) is hypothesised to promote localisation of this molecular Ca2+ dependent switch to synaptic regions and influence nearby downstream effectors of synaptic maintenance and plasticity. The long decay kinetics exhibited by GluN2B containing NMDAR can effectively integrate trains of stimuli by providing a long window for coincidence detection of synaptic release and post-synaptic depolarisation (Vargas-Caballero and Robinson, 2004) a function which may contribute to working memory in the prefrontal cortex where a substantial fraction of GluN2B-containing NMDARs remains in adulthood (Wang et al. 2008).

NMDARs are essential for numerous spatial memory tasks as well as for many forms of long-term synaptic plasticity, a molecular correlate of learning and memory (Bliss and Collingridge 1993; Takeuchi et al. 2013). Overexpression of the GluN2B subunit in mice led to increased recruitment of synaptic GluN2B, improved performance in memory tests and enhanced synaptic plasticity (Tang et al. 1999, Cui et al. 2011).

In adult and ageing rodents, the contribution of GluN2B to synaptic function is reduced compared to younger animals and the decline in GluN2B subunit expression is correlated with impaired memory functions (Clayton et al. 2002; Magnusson, 2012; Magnusson et al. 2010; Zhao et al. 2009). However, it was not known whether age dependent changes in GluN2B synaptic composition also occurred over the adult human lifespan. To analyse the age dependence of NMDAR composition in human synapses we studied samples from adult patients within a broad age span. Using GFAP and Iba1 as inflammation markers and GluN2B as a protein degradation marker, we were able to assess the quality of this tissue as non-pathological. We observed that although protein expression for both glutamate receptor subunits GluN2A and GluN2B is reduced with age, both proteins are still produced at detectable levels in older adults (Fig 2). By using a pharmacological approach, whereby Ro 25-6981 500 nM selectively inhibits receptors containing GluN1/GluN2B subunits, our data demonstrate that a significant fraction of these NMDAR channels participate in basal synaptic function in cortical slices in younger adults but less so in older adults. This result is in sharp contrast with predictions from equivalent mouse studies, and our own analyses where Ro 25-6981 had no significant effect of the NMDA/AMPA ratio in young adult mice (3-5 months) (Supplementary Figure 2 1A-C). Thus, our data using pharmacological blockade of synaptic currents with Ro 25-6981, and changes in τ_w_ in synaptic NMDAR currents shows that synaptic GluN2B contribution is reduced in human cortical synapses with ageing.

Mechanisms of synaptic anchoring via post-translational modifications (Lussier et al. 2015) and/or association with other synaptic density proteins (Chen et al. 2015) may underlie this age-dependent GluN2A or GluN2B content in human cortical synapses. Further analyses of human tissue derived from neurosurgery allowing for the analysis of well-preserved protein-protein interactions and post-translational modifications will allow a deeper understanding of the factors that cause GluN2B loss from older human synapses. Our measures do not distinguish between di-heteromeric receptors (GluN2B/GluN1) or tri-heteromeric receptors (GluN2B/GluN2A/GluN1), both of which show a widespread expression adult rodent brain (Rauner and Köhr, 2011) and further biochemical and/or pharmacological analysis will be required to address their contribution to human cortical synaptic transmission. Our measures focused on synaptic inputs to excitatory neurons, however, in rodents inhibitory interneurons also possess specific subsets of synaptic glutamatergic receptors including those containing GluN2B (reviewed in Akgül et al 2016) which are regulated in a regional and developmental manner. Further analysis will be required to test the synaptic composition in identified subtypes of human inhibitory interneurons.

Mounting evidence in model systems shows that a reduction in synapse number and turnover, as well as alterations of synaptic receptor composition, are correlated with age-related cognitive decline in rodents and primates (Morrison and Baxter, 2012; Mostany et al. 2013). Previous work suggests a causal role of GluN2B recruitment at the synapse in regulating synaptic strength and memory storage. Furthermore, overexpression of the GluN2B subunit in mice led to increased recruitment of synaptic GluN2B resulting in improved memory and enhanced synaptic plasticity (Cui et al. 2011; Tang et al. 1999). Since synaptic NMDAR content and composition can determine integrative and plastic properties in neurons (Yashiro and Philpot, 2009), our observations highlight a biologically plausible mechanism for the reduction in cortical plasticity observed across the human age span (Freitas et al. 2011) and the cognitive ageing in the human brain which can be observed from middle age (Singh-Manoux et al. 2012).

Experiments using live human tissue derived from neurosurgery can further our understanding of molecular mechanisms behind synaptic ageing synapse alterations in disease. This can be studied either directly in diseased tissue from patients (Finardi et al. 2006; Ying et al. 2004) or by acutely mimicking disease states using non-pathological tissue processed with the brain slice technique (Vargas-Caballero et al. 2016).

Understanding synaptic composition throughout the human age span can also inform drug development for neurological conditions affecting young or older individuals, as well as aid in understanding and ameliorating off-target effects of widely used drugs. As an example, fluoxetine (Prozac), also inhibits GluN2B-containing receptors (Kiss et al. 2012) and thus, may have differential side-effects in young and older brains. Furthermore, data on human synapse composition and function, such as our work presented here, can serve as a benchmark to further develop pluripotent stem cell models (Zhang et al. 2016) to mimic the GluN2A/ GluN2B synaptic composition for the disease model of interest.

## Funding

This work was supported by the Institute for Life Sciences, University of Southampton (Senior Research Fellowship to MVC) and Wessex Medical Trust (Innovation Grant to MVC). CMP was funded by the Gerald Kerkut Trust and the Institute for Life Sciences (PhD studentship).

## Acknowledgements

We thank Prof. Delphine Boche for helpful discussions on human brain tissue immunohistochemistry.

---

**Supplementary Figure 1.**
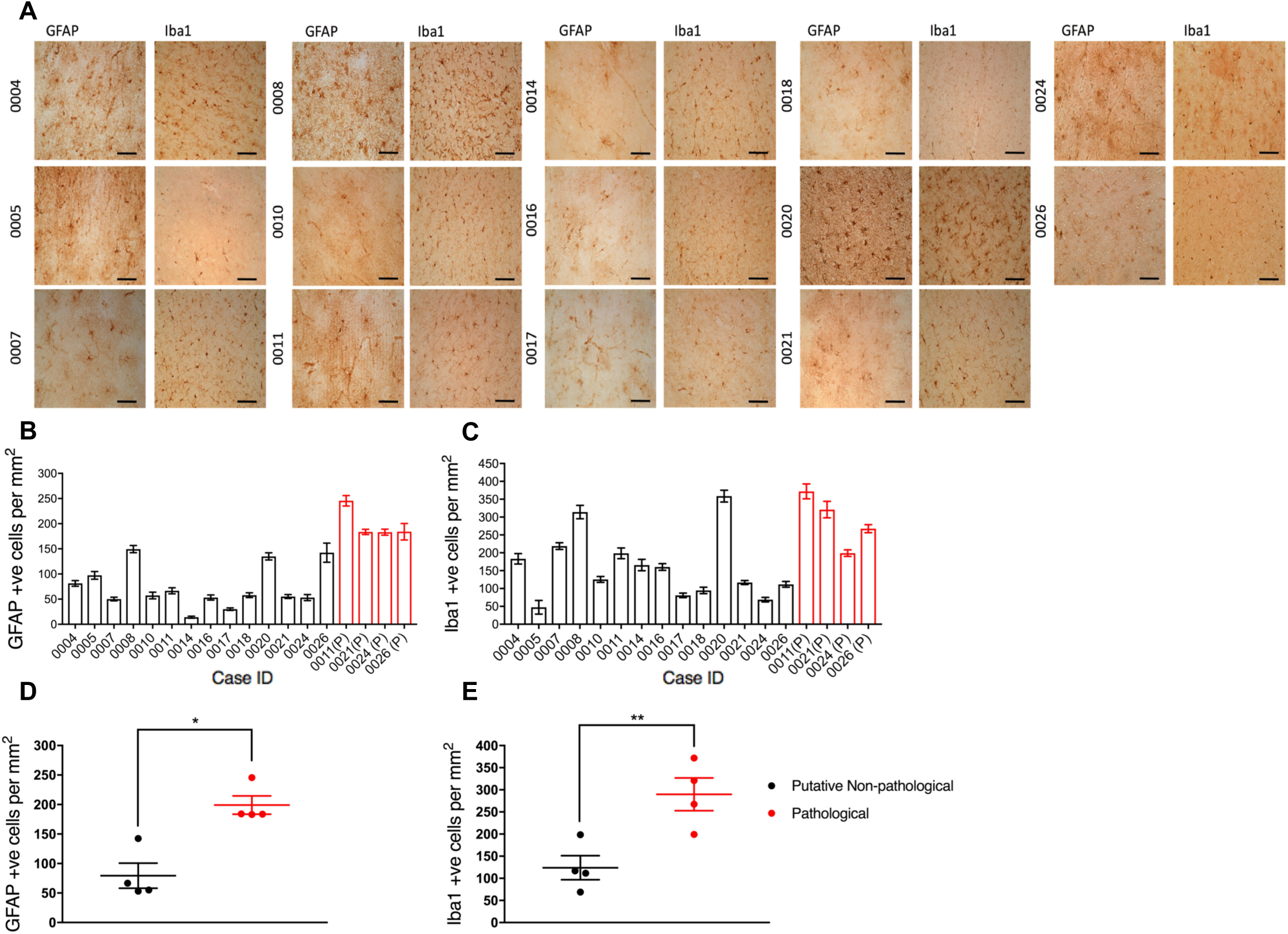
Representative images of immunohistochemical analyses for cases stained for GFAP and Iba1. (A). Cell density quantification for non-pathological (PNP) and pathological (P) tissue, for GFAP (B), and Iba1 (C). Data were analysed using one-way ANOVA with Dunnett’s post-hoc correction comparing against the mean of pathological cases. Cases 0008 and 0020 did not meet the criteria for subsequent analyses, as they were not significantly different to the mean of pathological cases (for Iba1). Paired comparison of cell densities for GFAP (D) and Iba1 (E) for putative non-pathological (PNP) tissue for which pathological (P) tissue was available (cases 0011, 0021, 0024 and 0026). Significantly lower densities for both GFAP and Iba1 positive cells revealed by paired t-tests (P = 0.0247 and P = 0.0018 respectively).

**Supplementary Figure 2.**
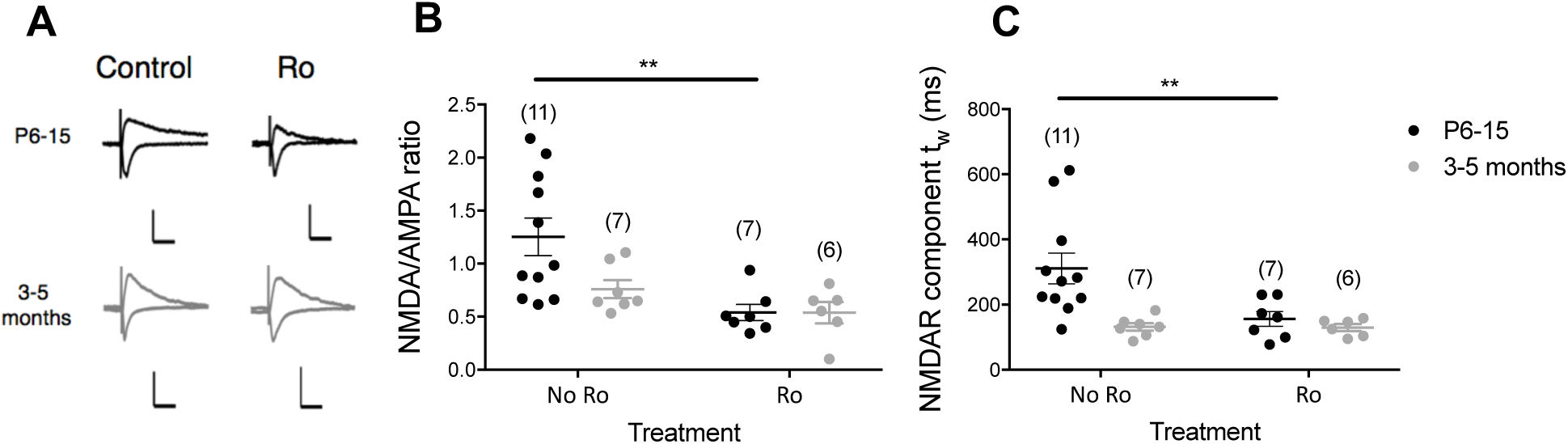
Significant GluN2B contribution in NMDA-mediated currents in temporal brain slices recorded from P6-15 mice but not from 3-5 month old (adult) mice. (A) Representative NMDAR- and AMPAR-mediated currents from P6-15 and 3-5 month-old mice with and without Ro 25-6981 treatment. (B) NMDA/AMPA ratios from P6-15 mice show sensitivity to Ro treatment, while recordings from 3-5-month-old mice do not. (C) Weighted NMDA time constant (τ_w_) shows sensitivity to Ro treatment in P6-15 mice. No differences were observed in the 3-5-month-old mice. Numbers in brackets correspond to number of cells recorded from a minimum of 4 mice. Scale bars in (**A**): 50ms, 100pA.

## Notes

#### Summary of Updates

minor changes

